# Using CarboTrace 480 to detect protoplastation in pigment deficient mutant of *Chlorella sorokiniana*

**DOI:** 10.64898/2026.08.27.747479

**Authors:** Sidsel Koggersbøl Thrane, Anders Olsen, Teis Esben Sondergaard

**Affiliations:** Department of Chemistry and Bioscience, Aalborg University, Fredrik Bajers Vej 7H, DK-9220, Aalborg, Denmark; Aliga ApS, Vandværksvej 12, DK-9800 Hjørring, Denmark

## Abstract

The increasing world population necessitates new sustainable nutrient sources, making microalgae like *Chlorella sorokiniana* interesting due to its rich nutrient profile and sustainable cultivation methods. With genetic optimization tools like CRISPR/Cas9, microalgae as a nutrient source can be improved even further. However, degradation of the rigid cell wall of microalgae, and thereby developing protoplasts, is often necessary prior to transformation, but monitoring protoplast development in spherical, single-celled organisms like *C. sorokiniana* is challenging using bright-field microscopy. Carbotrace 480 and 630 were tested as fluorescent markers of the cell wall of a *C. sorokiniana* mutant for protoplast detection, and Carbotrace 480 was successfully used to distinguish protoplast from normal cells in a cell suspension. The enzymes Driselase, Glucanex, Snailase, and Saczyme were tested in different combinations to degrade the cell wall of the mutant, with Snailase as the most effective yielding ∼60 % protoplasts. This study provides a quick and easy tool for monitoring protoplast development in the microalgae *C. sorokiniana*, the first step to improve *C. sorokiniana* as a sustainable nutrient source using genetic optimization tools like CRISPR/Cas9.

## 1. Introduction

Along with the increasing world population, the need for alternative and sustainable nutrient sources has increased. Traditional agriculture has a huge impact on the ecosystems, and thus it is not sustainable to meet the increase in the food production using only traditional agriculture. Instead, other nutrient sources like microalgae must be explored [1]. For this, microalgae such as *Chlorella*, have become interesting due to its broad nutrient composition, and the capability of being produced on non-arable land [2] and in closed systems, minimizing the impact on the environment [3]. The different microalgae-species can be protein-rich, containing all amino acids, it can be lipid-rich, containing the essential fatty acids EPA and DHA [4,5]. Although the microalgae can serve as a great source for sustainable nutrients in various applications, the genetic tools available today can improve this nutrient source even further.

With the genetic optimization tool CRISPR/Cas9, microalgae can be genetically optimized to increase e.g. the production of proteins, lipids etc, the productivity and yield during cultivation or other relevant aspects. The CRISPR/Cas9 system can be delivered to the cell through various methods, e.g. electroporation and PEG mediated transformation, but due to the cell wall, removing the cell wall, and thereby generating protoplasts, prior to transformation is necessary [6]. The cell walls of algae are in general recalcitrant to mechanical and chemical degradation, due to the complex structure [7]. The cell wall of *Chlorella* species consists of a complex single-layer cell wall with chitin- or chitosan-like fibers, which is embedded in a hemicellulose matrix, composed of of galactose, xylose, mannose, rhamnose and glucosamine, as well as amino sugars and uronic acids and the right combination of different enzymes is needed to degrade it [8].

When developing protoplasts of multicellular organisms with adhering cells, such as plant tissue or mycelium from fungi, a simple examination of the cell morphology in the culture will reveal if protoplasts are starting to form. Spherical singled cells will start to form, indicating that the cell wall degrading treatment is working [9]. But when developing protoplasts from *Chlorella* species or similar organisms, it can be very difficult to distinguish normal cells from protoplasts, since the appearance of the protoplasts will not be clearly distinct from normal cells. Studies have described how the size of the protoplasts may change due to the osmolarity of the buffer solution [10–12]. Because cell sizes already vary several µm within the same suspension prior to enzyme treatment, due to the cells being in different stages of cell division [13], it is difficult to rely on size alone as an indicator of protoplast formation. This becomes particularly problematic when the enzyme treatment yields only a low proportion of protoplasts, as is often the case before proper optimisation. Therefore, a simple and reliable method to distinguish intact cells from protoplasts is needed.

One approach is to stain the cell wall with a fluorescent dye and compare the same cell suspension using both bright-field and fluorescence microscopy. Several fluorescent markers are commercially available for detecting cell walls in microalgae, including traditional stains such as Calcofluor White [14] and more recent alternatives like CarboTrace, an optotracer that binds to the glycosidic linkages in cellulose [15]. However, there is still limited information on how CarboTrace interacts with microalgal cell walls. This study therefore explores its applicability for protoplast detection in Chlorella sorokiniana.

## 2. Materials and methods

### 2.1 Strains and culture conditions

The experiments were conducted on a pigment deficient mutant of *Chlorella sorokiniana* (M12) (Thrane et al. 2023). The strain was cultivated in liquid heterotrophic medium [17] in dark at 32°C, 150 RPM.

### 2.2 pH optimum

Different enzymes have different pH optima, but the selected pH must also support the viability of the *Chlorella sorokiniana* culture during protoplast formation. An experiment was therefore conducted to determine which pH levels, within the range of 3–10, could sustain culture growth. The pH of the liquid heterotrophic medium was adjusted using either HCl or NaOH and subsequently sterilised using a 0.22 µm filter. The medium was inoculated with biomass from M12 and incubated for 72 h. OD_750 nm_ was measured 2 hours after inoculation and three times during the incubation period. The experiment was done in biological triplicates.

### 2.3 Protoplast preparation

The enzymes Driselase (Sigma-Aldrich), Glucanex (Sigma-Aldrich), Snailase (Abbexa) and Saczyme (Novonesis) were used in different combinations to evaluate cell wall degradation. Biomass was harvested by centrifugation at 2000 x *g* at RT for 2 min and the supernatant was discarded. Cells were washed by resuspending pellets in Protoplast Buffer Solution comprising 0.6 M D-Mannitol and 25 mM Tris (pH adjusted to 6 w. HCl/NaOH), following centrifugation at 2000 x *g* at RT for 2 min and supernatant was discarded. The harvested and washed biomass was resuspended in Protoplast Buffer Solution also comprising one of the following enzyme combinations: ① 1% Driselase + 1% Glucanex, ② 1% Driselase + 1% Glucanex + 1% Snailase, ③ 1% Driselase + 1% Glucanex + 1% Saczyme, ④ 1% Snailase and ⑤ 1% Saczyme. Due to the un-dissolvability of Driselase, the enzyme mix was prepared as described by [18]. The cultures were incubated at 32°C, 100 rpm, in the dark for 21 hours with gentle inversion on HulaMixer.

### 2.4 Staining with Carbotrace 480 and 630

Undigested cells were stained with CarboTrace 480 and 630 (Ebba Biotech AB, Stockholm, Sweden) to determine which dye was most suitable for detecting protoplast formation. Each dye was diluted 1:3000 in phosphate-buffered saline (PBS, pH 7.4) for working solutions. Aliquots of 500 µL culture were centrifuged at 2000 × g for 2 min at room temperature (RT), after which the supernatant was discarded, and the pellet resuspended in 50 µL of staining solution. The samples were incubated for 30 min at RT with gentle mixing by inversion twice during the incubation period. Following incubation, cells were washed by centrifugation (2000 × g, 2 min, RT), the supernatant discarded, and the pellet resuspended in 500 µL PBS; this washing step was repeated to remove unbound dye. Finally, 5 µL of the stained suspension was mounted on microscope slides and imaged using an Olympus IX83 inverted confocal microscope (Evident Corporation, Tokyo, Japan) with a Yokashawa CSU-W1 spinning disk unit (Yokashawa Electric Corporation, Nakacho, Japan). CarboTrace 480 was excited using a 405 nm laser and emission detected with a B525/50 filter, whereas CarboTrace 630 was excited with a 488 nm laser and emission detected using a B617/73 filter.

To assess the enzymatic degradation of the cell wall in M12, cultures were stained with CarboTrace 480. Aliquots of 500 µL from each protoplast suspension were centrifuged at 2000 × g for 2 min at RT, after which the supernatant was discarded, and the pellet gently resuspended in 1 mL PBS. Cells were then stained as described above using CarboTrace 480 with one modification; all resuspension steps were performed with particular care to avoid rupturing potential fragile protoplasts. Following staining, 5 µL of the suspension was mounted on microscope slides and examined by both bright - field and fluorescence microscopy at 400× magnification, on a Leica DMI3000 B inverted microscope (Leica Microsystems GmbH, Wetzlar, Germany) using pE-300^white^ LED (CoolLED, Andover, England) for excitation at 450 nm and a 525/50 filter set for detection of emission. Images were acquired with both bright-field and fluorescence microscopy at identical focal planes. Cells visible in bright-field versus fluorescence were manually counted for each enzyme combination to assess the extent of cell wall degradation. For each sample, images were analysed at three independent focal planes.

## 3. Results

Cultivation across a pH range of 3–10 showed that the M12 mutant was able to grow between pH 6 and 10, although growth at pH 10 was limited. In contrast, no increase in OD_750_ was observed at pH 3–5, indicating that cells were either non-viable or not actively growing (Figure 1).

**Figure 1.**
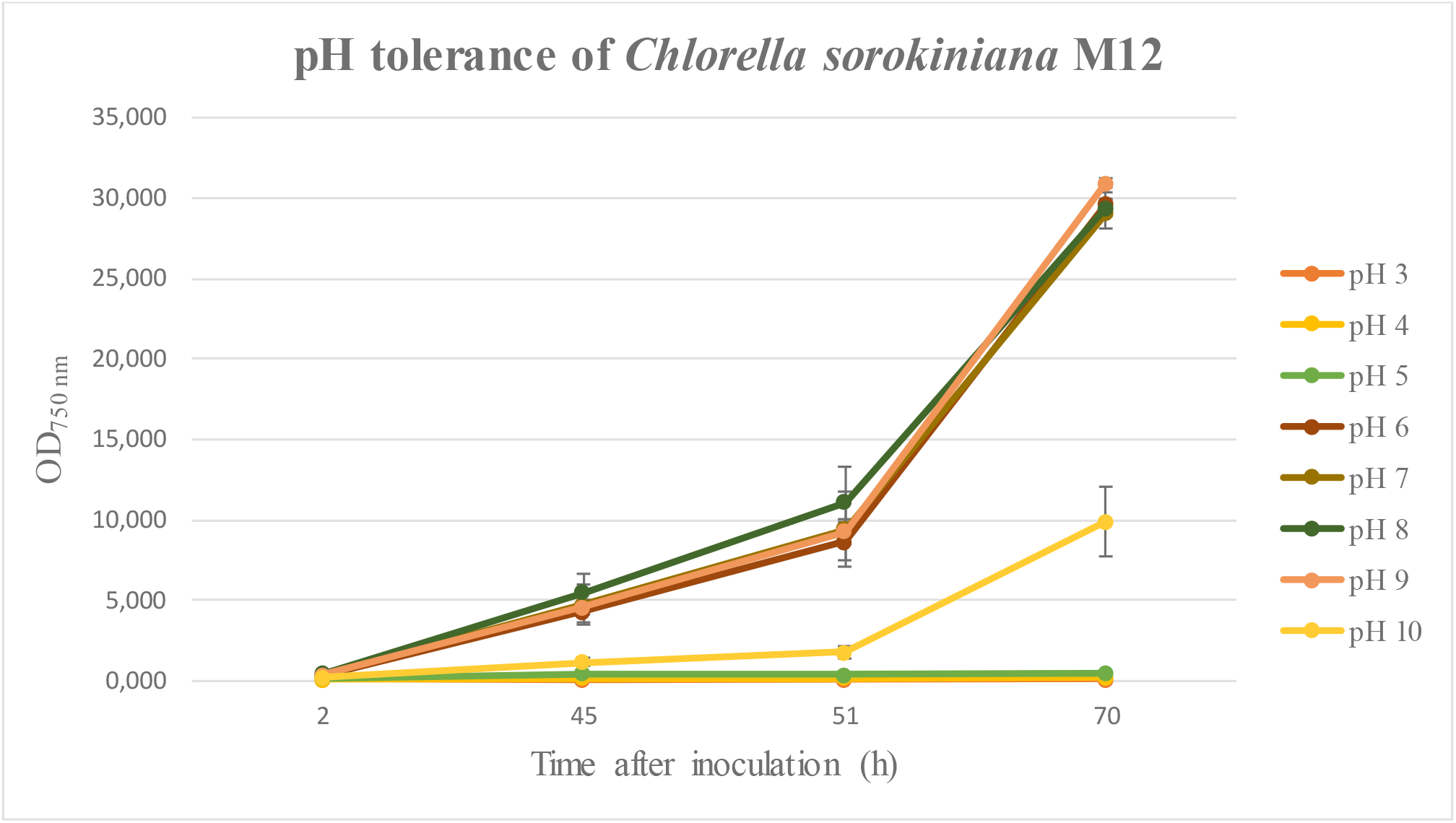
OD_750 nm_ of M12 cultivated at different pH levels. Cultures at pH 6-9 yielded similar growth, at pH 10, the cultures were growing but limited, and at pH 3-5 the cultures were not growing.

Image from spinning disc confocal microscopy (Figure 2) shows Carbotrace 480 and 630 are both binding to the cell wall. The image also shows that when using Carbotrace 630, there is a fluorescent signal from the interior of the cell (Figure 2, B), whereas there is no fluorescence from intracellular structures, when using Carbotrace 480 (Figure 2, A).

**Figure 2.**
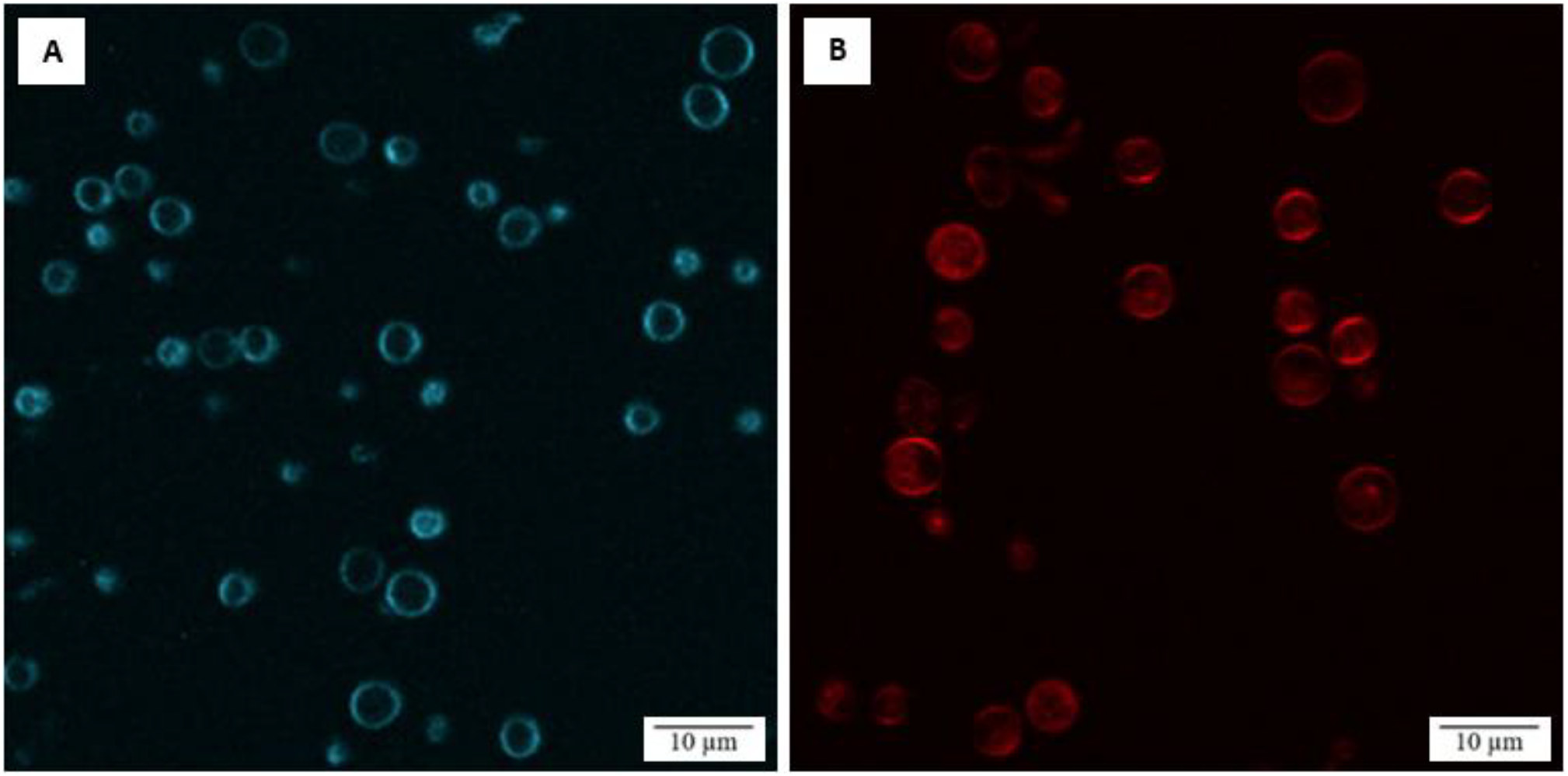
Spinning disc confocal microscopy of *Chlorella sorokiniana* mutant stained w. A) Carbotrace 480 and B) Carbotrace 630.

Carbotrace 480 was used to stain the cell wall of M12 after digestion with different enzyme combinations, to evaluate the effect of the enzyme mix on the cell wall to develop protoplasts. After enzyme digestion and staining, images of the cultures were acquired using bright-field and fluorescence microscopy at the same focal plane (Figure 3) Cells visible with both imaging techniques were considered to have intact cell walls (C), whereas cells observed only in bright-field microscopy were classified as protoplasts (P). Some cells were detectable with both imaging techniques but exhibited weak fluorescence, which were most likely cells with partially degraded cell walls.

**Figure 3.**
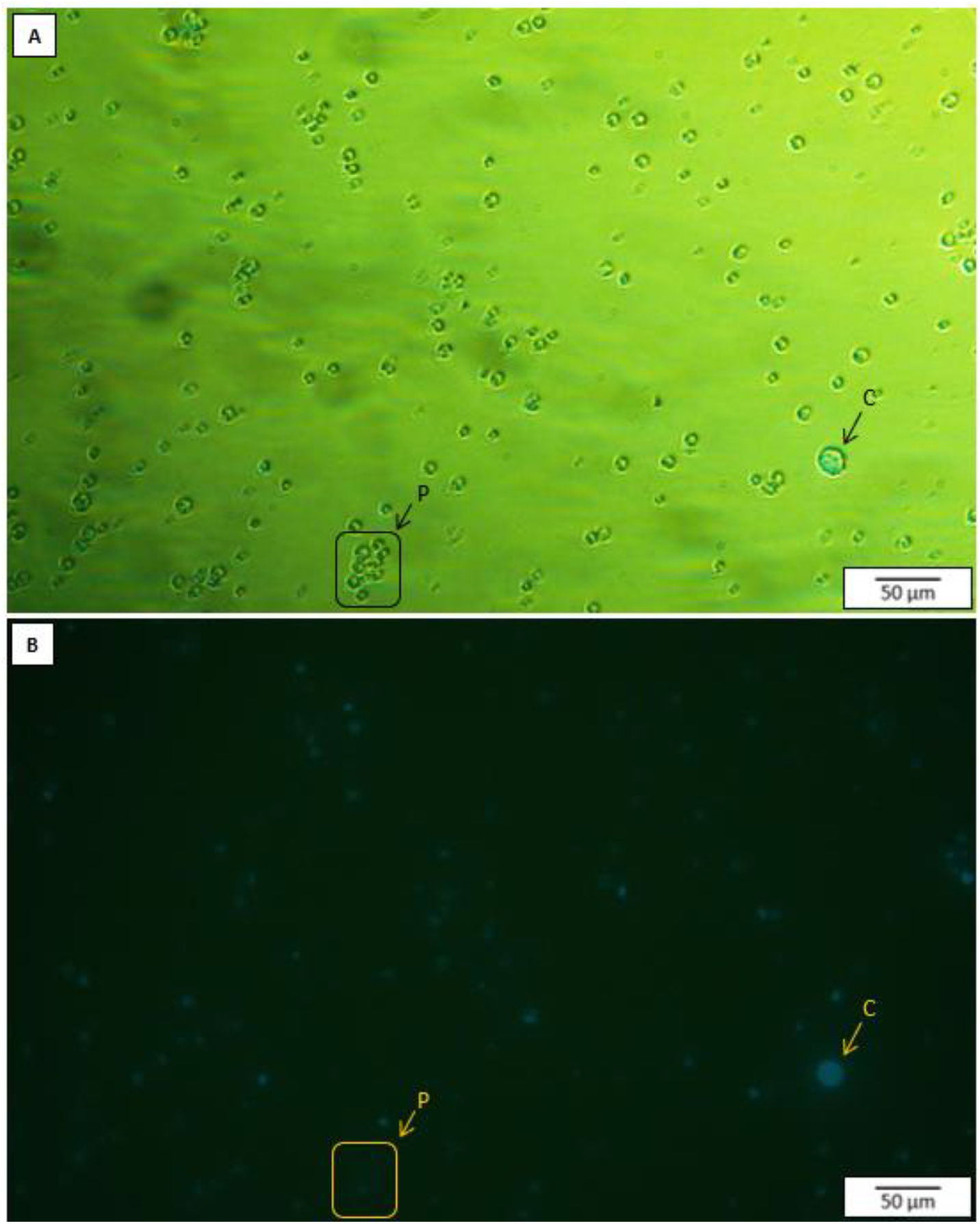
Pictures of *Chlorella sorokiniana* mutant stained with Carbotrace 480 after digestion with 1% Snailase. A) All cells are visible with brightfield microscopy. B) Only cells stained with Carbotrace are visible, using fluorescence microscopy. C marks an example of a normal cell with a cell wall that has been stained with Carbotrace 480. P marks protoplasts, cells that are only visible with brightfield microscopy, since the cell wall has been degraded, and the cell is therefore not stained.

Some enzyme combinations yielded protoplasts, enzyme mix nr. 4 (1% snailase) yielded ∼60% protoplasts as the most effective and enzyme mix 1 (1% Driselase and 1% Glucanex) as well as enzyme mix 3 (1% Driselase, 1% Glucanex and 1% Saczyme) both yielded ∼20% (Figure 4, B).

**Figure 4.**
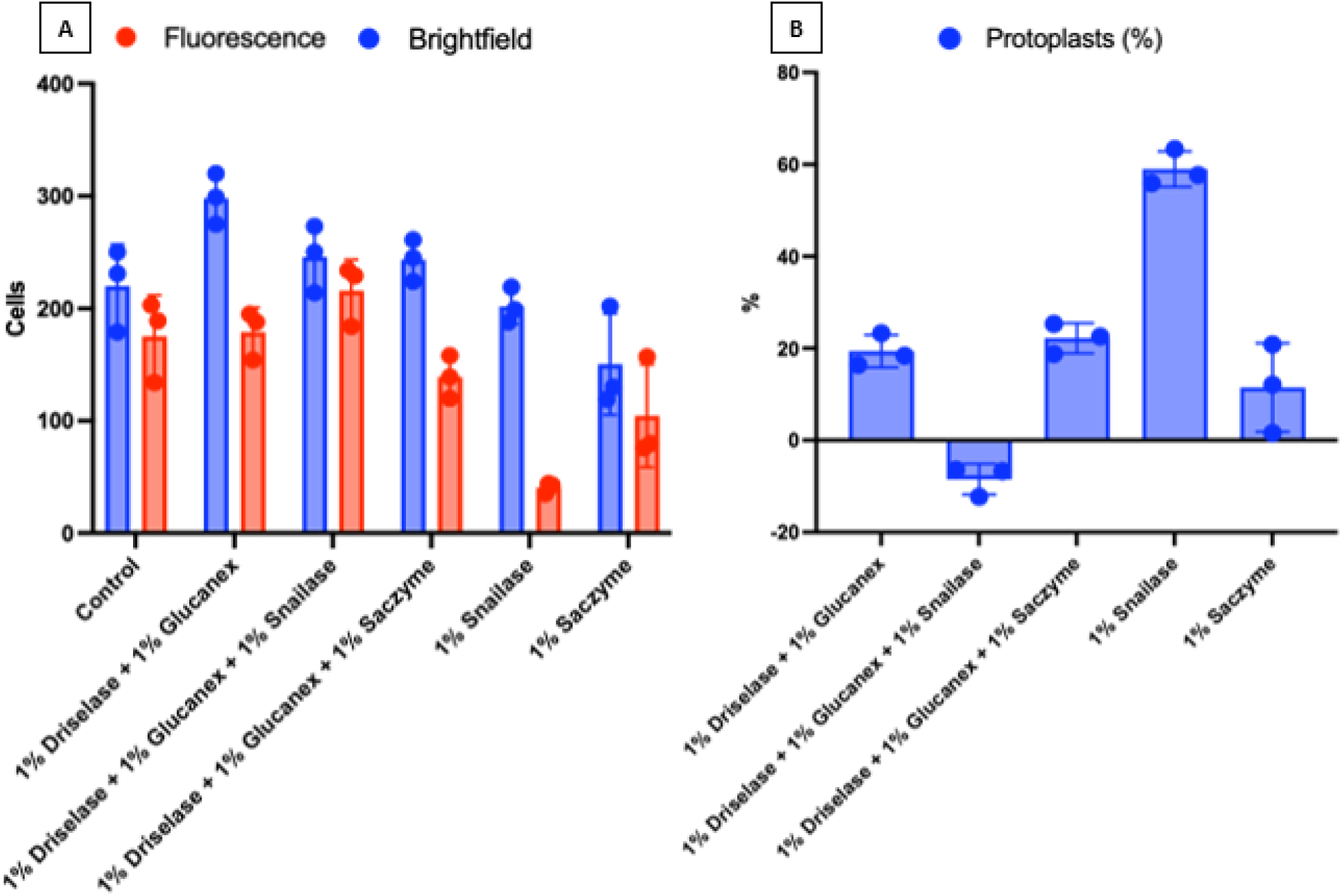
A) Number of cells visible with bright-field microscopy (blue) and fluorescence (red) after treatment with different enzyme combinations and B) Percentage of protoplasts developed with the different enzyme combinations.

## 4. Discussion

Using spinning disk confocal microscopy, it was confirmed that CarboTrace 480 binds specifically to the cell wall of the *Chlorella sorokiniana* mutant (M12), with no detectable staining of intracellular structures. This indicates that internal components, which remain present in protoplasts, do not contribute to the fluorescent signal. The images further demonstrate the absence of autofluorescence from intracellular structures at used filtersettings. In contrast, staining with CarboTrace 630 also labelled the cell wall but revealed additional fluorescence from intracellular regions, most likely due to chlorophyll autofluorescence in the chloroplasts. Chlorophyll *a* and *b* each exhibit two excitation peaks, with the first around 430 and 453 nm, respectively [19], but can also be excited by the 488 nm laser line used for CarboTrace 630. Their emission peaks, around 662 and 642 nm [19], overlap with the detection range of the B617/73 filter used for CarboTrace 630. Consequently, CarboTrace 480 is more suitable for protoplast detection, as the fluorescence signal is restricted to the stained cell wall.

Identifying an effective combination of cell wall–degrading enzymes is often one of the most time-consuming steps in protoplast generation, since cell wall composition varies between species and is often poorly characterized, and optimisation is therefore typically required. Establishing an efficient enzyme mixture early can therefore significantly reduce experimental time and improve overall yield and reproducibility. CarboTrace 480 was used to monitor protoplast formation following digestion with different enzyme combinations. The cell wall of *Chlorella sorokiniana* consists of multiple components, and as for many species within the Chlorophyta division, its detailed composition and structure remain incompletely characterised. Five enzyme combinations were therefore tested to evaluate their efficiency in degrading the cell wall. Saczyme alone did not produce a measurable effect, and its inclusion in combinations with Driselase and Glucanex did not improve protoplast formation. This is likely due to its primary activity on complex carbohydrates, whereas Glucanex contains activities such as chitinase that target chitin-like components of the cell wall. The most effective enzyme for cell wall degradation was Snailase, a mixture of enzymes derived from the digestive tracts of snails, which are natural predators of microalgae [20,21]. However, combining Snailase with Driselase and Glucanex did not enhance cell wall digestion, suggesting that excessive enzyme complexity may lead to inhibition or interference between enzymatic activities. Although treatment with 1% Snailase resulted in approximately 60% protoplast formation, further optimisation of enzyme concentration and incubation time could likely improve yields.

## 5. Conclusion

This study tested two types of cell wall stains, Carbotrace 480 and 630, and four cell wall degrading enzymes, Driselase, Glucanex, Snailase and Saczyme, to be used for protoplast development and detection of a mutant of the microalgae *Chlorella sorokiniana*. Carbotrace 480 was successfully used to detect protoplast in a cell suspension. Out of five different enzyme combinations, 1% Snailase was the most effective, yielding ∼ 60% protoplasts. This study introduces a method for detecting protoplast development, a process difficult to monitor in cultures of the spherical, single-celled organism *C. sorokiniana*, as well as adding to the knowledge of cell wall degrading enzymes effect on the cell wall of *C. sorokiniana*.

